# Cobamide-Dependent Dichloromethane Fermentation by *Dehalobacter* Reveals a Hidden Acetogenic Route for Organohalide Biotransformation

**DOI:** 10.64898/2026.05.26.727916

**Authors:** Huijuan Jin, Xiuying Li, Xin Wang, Hongyan Wang, Jingjing Wang, Ke Shi, Guanchi Liu, Tongyue Zhou, Siqi Huang, Michael Manefield, Frank E. Löffler, Jun Yan, Yi Yang

## Abstract

Halogenated one-carbon (C1) compounds, such as dichloromethane (DCM), drive critical fluxes in global carbon and halogen cycles. While the genus *Dehalobacter* is canonically defined by obligate organohalide respiration, its physiological and ecological roles in anaerobic C1 metabolism have remained fundamentally ambiguous. Here, we document a paradigm-shifting metabolic capacity within a sediment-derived microbial consortium: the autonomous fermentation of DCM by a novel population, ‘*Candidatus* Dehalobacter formatiformans’ strain J1. Over successive transfers, strain J1 outcompeted co-existing *Dehalobacterium formicoaceticum* to become the overwhelmingly dominant population (>80% relative abundance), converting DCM stoichiometrically to acetate and formate (4:1) without auxiliary substrates. Genome-resolved metagenomics revealed that strain J1 couples a distinct *mec* gene cassette—mediating methyl-transfer reactions during DCM activation—to a complete Wood-Ljungdahl pathway for efficient C1 assimilation. Crucially, strain J1 lacks the complete genetic repertoire for *de novo* cobamide biosynthesis. Physiological validation confirmed that this fermentative pathway is strictly dependent on exogenous cobamides, exposing a profound reliance on community cross-feeding. These findings reveal an unexpected acetogenic lifestyle within *Dehalobacter*, a lineage historically viewed as comprising obligate organohalide-respiring bacteria. More broadly, this work identifies cobamide-dependent methyl-transfer metabolism as an ecological control on anaerobic DCM fermentation and expands the known roles of *Dehalobacter* in carbon–halogen cycling in anoxic environments.

## Introduction

Dichloromethane (DCM), is a prevalent halogenated one-carbon (C1) compound ubiquitously distributed across atmospheric, aquatic, and sedimentary environments, where it plays a significant role in the global carbon and halogen cycling as both a source and sink of reactive carbon and chlorine species [1, 2]. Although natural sources including wetlands, oceans, geothermal activity, and biomass burning, contribute approximately 80,000 tons annually, anthropogenic emissions predominate due to DCM’s widespread industrial use in pharmaceuticals, plastics manufacturing, paint stripping, adhesive production, and related applications[3–6]. Global DCM production has continued to rise and is now dominated by the Asia-Pacific region driven by industrial and pharmaceutical demand, with China alone emitting approximately 628 Gg y⁻¹ in 2019 [6]. Improper disposal and prolonged exposure to DCM have been associated with adverse health effects, including an elevated risk of liver cancer [7–9]. Accordingly, the International Agency for Research on Cancer (IARC) has classified DCM as a Group 2A probable human carcinogen (https://monographs.iarc.who.int/). Owing to its well-documented toxicity and exposure risks, DCM is now subject to increasingly stringent frameworks worldwide, including usage bans and restrictions enforced by China’s Ministry of Ecology and Environment (MEE), and high-priority hazardous listings by the Agency for Toxic Substances and Disease Registry (ATSDR).

Given these challenges, increasing attention has been directed toward microbial transformation pathways capable of degrading DCM. Recent studies have highlighted the ecological and biogeochemical importance of DCM turnover, revealing that its microbial degradation pathways contribute not only to halogen detoxification but also to carbon fluxes in various ecosystems [10]. Certain microorganisms can utilize DCM as carbon and energy source, thereby mediating its bioconversion into cellular constituents or inorganic products, and contributing to the environmental cycling of carbon and chlorine. Aerobic bacteria, including species from genera such as *Albibacter* [11], *Bacillus* [12], *Ancylobacter* [13], *Methylobacterium* [14], *Hyphomicrobium* [15]*, Methylophilus* [14]*, Lysinobacillus* [16], *Gottschalkia* [17] *, Xanthobacter* [18], *Paracoccus* [19] and *Methylopila* [20], are capable of using DCM as a sole source of carbon and electrons, expressing a conserved glutathione-dependent dehalogenase (DcmA) that catalyze DCM into formaldehyde and two HCl molecules [21–23], and then to downstream C_1_ assimilation pathways such as the serine or ribulose monophosphate (RuMP) cycles [24]. Under anoxic conditions, DCM is often readily degraded by diverse *Peptococcaceae* species though generally conserved metabolic pathways. Among these, *Dehalobacterium formicoaceticum* DMC is the only isolate described that can anaerobically metabolize DCM to acetate and formate (1:2 molar ratio) [25–28], while ‘*Candidatus* Formimonas warabiya’ produces only acetate and can also utilize non-chlorinated substrates like methanol, choline, and N,N,N-trimethylglycine [29, 30]. ‘*Candidatus*

Dichloromethanomonas elyunquensis’ consortium RM is a more recent discovery enriched from pristine river sands in Puerto Rico, which involves the complete mineralization of DCM to H_2_ and CO_2_ [31–34]. These *Peptococcaceae* members share a complete Wood-Ljungdahl pathway (WLP) and encode a conserved DCM catabolism (*mec*) gene cluster (8-10 genes), which includes multiple methyltransferases [10]. Current evidence suggests that the *mec* cassette mediates the initial activation and dechlorination of DCM, transferring the methylene carbon to tetrahydrofolate as a C1 unit that subsequently enters the WLP for fermentation, acetogenesis, or complete oxidation. Metaproteomic analyses confirm high expression of the *mec* cassette in these DCM-degrading cultures, underscoring its central role in anaerobic DCM catabolism [35, 36].

Several studies have reported that *Dehalobacter* can persist and potentially grow in DCM-amended mixed cultures, with acetate and/or CO_2_ as major end products [37–41]. However, for years, the functional attribution of this activity to *Dehalobacter* remained uncertain, partly due to taxonomic ambiguity. For example, one population initially interpreted as a DCM-degrading Dehalobacter from consortium RM was later reclassified through genome-resolved phylogenetic analysis [31]. In a separate enrichment, a *Dehalobacter*-affiliated phylotype was outcompeted by “*Ca. Formimonas warabiya*” during DCM incubation [29]. These observations have complicated the functional attribution of DCM degradation to *Dehalobacter* and reinforced the view that, until recently, members of this genus were primarily recognized as obligate organohalide-respiring bacteria rather than bona fide DCM-fermenting microorganisms. Only recently has multi-omics evidence provided strong support for *Dehalobacter*-mediated DCM degradation, indicating that the direct involvement of this genus in anaerobic DCM turnover is an emerging discovery [38, 41]. In the DCM-degrading microbial consortium DCME, which is dominated by ‘*Ca.* Dehalobacter alkaniphilus’ strain DAD, methane is produced as the major end product. This observation, similar to findings from consortium RM, has been interpreted as evidence for anaerobic DCM mineralization [38, 42, 43]. However, direct physiological evidence for DCM-dependent growth by strain DAD, as well as genomic evidence for *mec*-mediated DCM activation by this organism, remains limited. Thus, the precise metabolic role of strain DAD in DCM transformation has not yet been fully resolved.

In this study, we investigated anaerobic DCM degradation in a stable, non-methanogenic enrichment culture in which two putative DCM-degrading populations, *Dehalobacterium* and *Dehalobacter*, were consistently detected. In successive subcultures, *Dehalobacter* became the overwhelmingly dominant DCM degrader, suggesting a competitive advantage over *Dehalobacterium* under nutrient-limited conditions. Through integrated physiological and genomic analyses, we identified a novel *Dehalobacter* lineage that mediates anaerobic fermentative DCM degradation and possesses a distinct complement of conserved and divergent genes involved in DCM metabolism. These findings expand current knowledge of DCM degradation by *Dehalobacter*, demonstrating that it can transform DCM via a non-mineralizing fermentative pathway and refining its previously assumed metabolic role in anaerobic DCM turnover, with implications for chlorinated C_1_ compound cycling in oligotrophic subsurface environments.

## Results

### Anaerobic DCM degradation in microcosms and transfer cultures

During a two-week incubation, the initial dose of DCM (154.9 ± 2.8 μmoles) was completely consumed in the live microcosms (Figure 1A), whereas DCM consumption was negligible in autoclaved controls (Figure 1B). After three successive transfers in lactate-amended medium, sediment-free enrichment cultures were obtained that degraded 161.8 ± 2.1 μmoles of DCM at an average rate of 323.5 ± 4.2 μM per day. The DCM degradation was accompanied by the formation of acetate (148.1 ± 9.0 μmoles), propionate (261.2 ± 15.0 μmoles), and formate (6.8 ± 0.9 μmoles) (Figure 1C). DCM conversion rates increased with consecutive transfers, reaching 372.2 ± 6.3 μM per day in the 10^th^ transfer cultures (Figure 1D). In subsequent transfers without lactate, propionate production ceased, while DCM degradation and formation of acetate and formate persisted. In the 11^th^ lactate-free transfer, 156.6 ± 2.9 μmoles DCM was consumed at a rate of 229.4 ± 4.0 μM d⁻¹, yielding 39.1 ± 8.1 μmol acetate and 10.5 ± 1.7 μmol formate after 7-day (Figure 1E), indicating that lactate was not essential for DCM degradation. Chloromethane (CM) and methane (CH_4_) were not detected in any microcosms or transfer cultures, suggesting that DCM was not degraded via reductive dechlorination and that methanogenesis was completely suppressed. By the 15^th^ lactate-free transfer, DCM degradation rates (410.9 ± 9.7 μM d^-1^) were comparable to those observed in lactate-amended transfers. The terminal products were acetate (44.4 ± 4.4 μmol) and formate (9.8 ± 1.4 μmol), consistently exhibiting an approximate molar ratio of 4:1 (Figure 1F). Collectively, these results indicated that microorganism(s) originating from Xi River sediment are capable of anaerobic DCM metabolism, leading to the production of acetate and formate by a non-mineralizing fermentative pathway.

**Figure 1.**
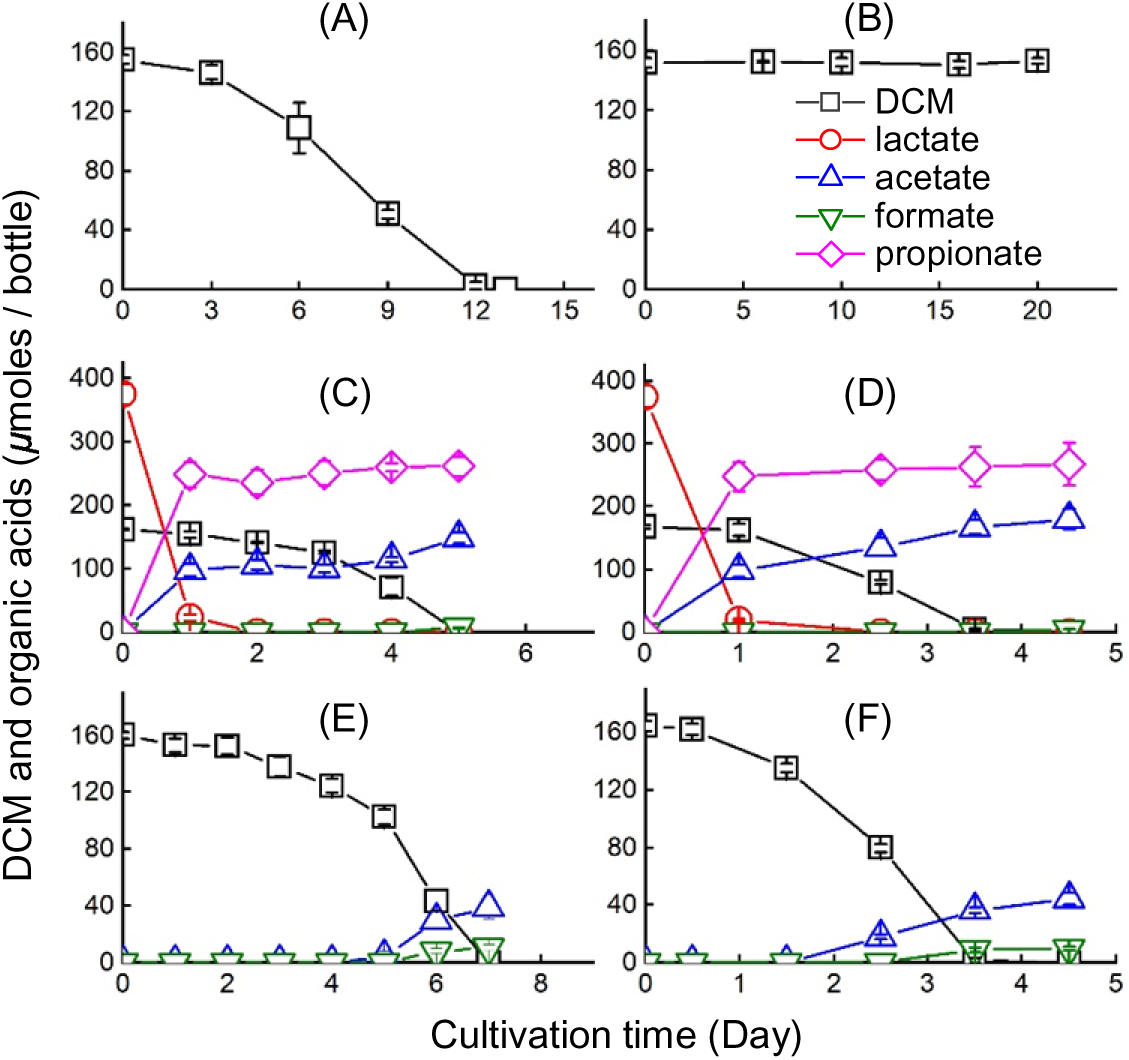
Anaerobic degradation of DCM and formation of transformation products during microcosm incubation and successive transfers. (A) DCM consumption in live microcosms. (B) DCM concentrations in autoclaved controls demonstrating negligible abiotic loss. (C) DCM degradation and metabolite production (acetate, propionate, and formate) in the third transfer culture amended with lactate. (D) DCM degradation and metabolite production in the tenth transfer amended with lactate. (E) DCM degradation and transformation in the eleventh transfer. (F) DCM degradation and transformation in the fifth transfer. Error bars represent the standard deviations of three biological replicates.

### Microbial community analysis of the DCM-degrading enrichment cultures

The microbial diversity of the enrichment culture decreased markedly with successive transfers. At the genus level, the dominant bacterial taxa in the DCM-fed transfer cultures amended with lactate included *Clostridium* sensu stricto 7, *Dehalobacter*, *Sedimentibacter*, *Christensenellaceae* R-7 group, *Lentimicrobium*, *Anaerostigum*, and *Paludibacter*. Among these genera, *Dehalobacter* exhibited a pronounced increase in relative abundance, rising from 5.4% in the 3^rd^ transfer to 59.9% in the 10^th^ transfer. In contrast, *Clostridium* sensu stricto 7, *Sedimentibacter*, and *Anaerotignum* each accounted for less than 22.0% of the total community (Figure 2). Upon removal of lactate, *Dehalobacter* became the most abundant organism, representing 76.3% and 81.4% of the community in the 11^th^ and 15^th^ transfer cultures, respectively. Notably, this dominance was maintained despite the absence of an external electron, while *Sedimentibacter* persisted at lower abundances (8.7-11.7%). Other genera, including *Christensenellaceae* R-7 group, *Monoglobus*, unclassified *Rikenellaceae*, *Proteiniclasticum*, and *Desulfomicrobium*, were detected only at trace levels (< 1%). Low-abundance populations affiliated with ‘*Ca.* Dichloromethanomonas elyunquensis’ were also detected (0.04% and 0.02% in the 11^th^ and 15^th^ transfers, respectively) (Figure 2). The consistent enrichment and dominance of under DCM-fed, lactate-free conditions suggest that this organism is capable of directly utilizing DCM as an energy and carbon source, supporting its role as a key DCM degrader in this system implicating *Dehalobacter* as a potential anaerobic fermentative DCM-metabolizing bacterium.

**Figure 2.**
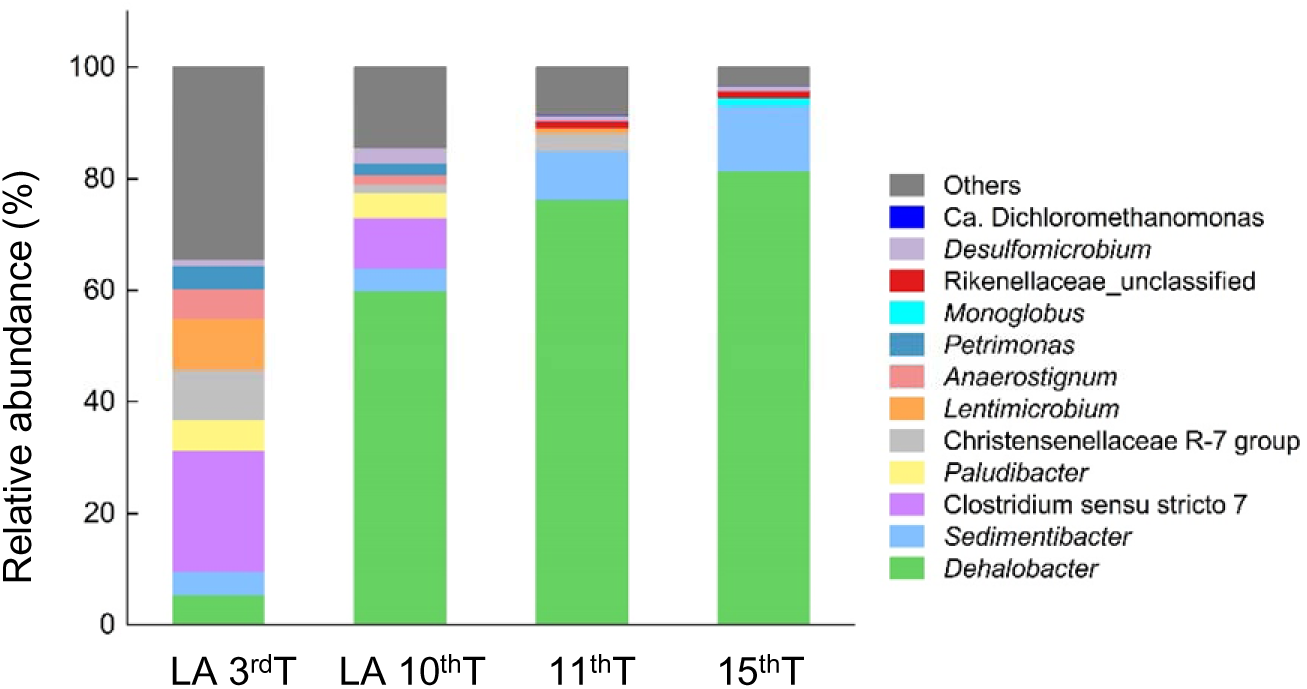
Bacterial community composition across different transfers. LA indicates supplementation with 10 mM lactate. 3^rd^ T, 10^th^ T, 11^th^ T and 15^th^ T denote the third, tenth, eleventh and fifteenth transfer cultures, respectively.

### Metagenome comparative analyses

Based on taxonomic assignments against the NCBI-nr database, sequences affiliated with the genera *Dehalobacter* (19.6%) and *Dehalobacterium* (14.2%) were most abundant, followed by *Lentimicrobium* (9.1%), *Clostridium* (4.2%), *Desulfosporosinus* (4.2%), ‘*Ca.* Dichloromethanomonas elyunquensis’ (1.8%), and *Anaerotignum* (1.7%) (Figure 3A). Notably, *Dehalobacterium* was not detected in the amplicon sequencing dataset (Figure 2), but was recovered through metagenomic sequencing, highlighting discrepancies between taxonomic profiles derived from amplicon-based and shotgun approaches. In total, 11,647 contigs were assembled, with an *N_50_* of 16,017 bp and a maximum contig length of 768,654 bp. Following assembly and binning of quality-filtered metagenomic reads, 12 high-quality metagenome-assembled genomes (MAGs) were recovered, each exhibiting >70% completeness and <10% contamination (Table S1). Six MAGs were classified at the genus level, including *Dehalobacter* (bin_3), *Propionicimonas* (bin_6), *Lentimicrobium* (bin_7), *Petrimonas* (bin_8), *Metalachnospira* (bin_12) and Bact-08 (bin_13). Two MAGs were identified at the species level as *Anaerotignum propionicum* (bin_05) and *Dehalobacterium formicoaceticum* (bin_11). Additionally, three MAGs were assigned to the family level, namely *Clostridiaceae* (bin_1), UBA5745 (bin_2), and *Oscillospiraceae* (bin_4), and one MAG was class *Negativicutes* (bin_9) (Table S1).

**Figure 3.**
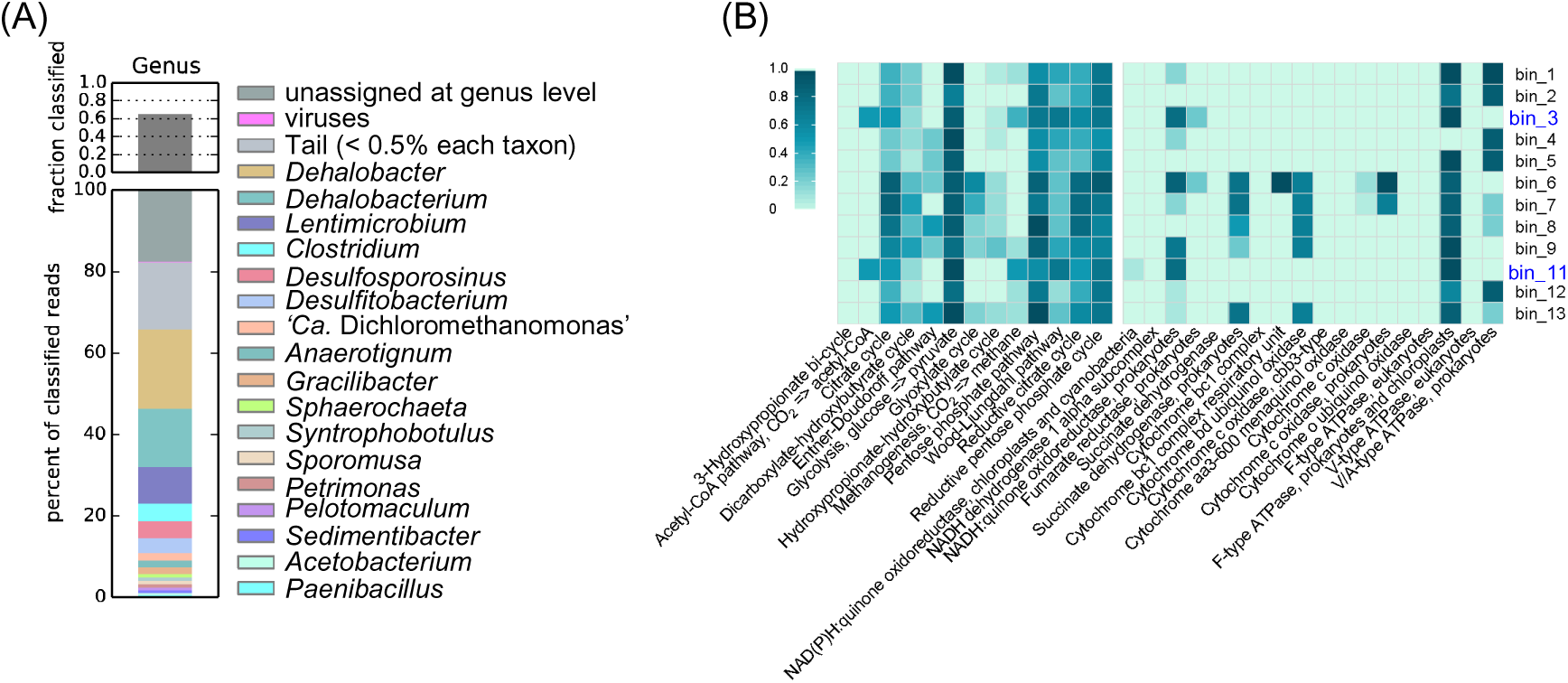
Taxonomic composition and metabolic potential of metagenomic assemblies. (A) taxonomic classification of shotgun metagenomic read data with Kaiju v1.7.3. (B) Metabolic pathway reconstruction and subunit completeness of MAGs based on DRAM annotation profiles. Pathways, subunits, or process absent from all high-quality bins were omitted for clarity. Bin.3 and Bin.11, representing *Dehalobacter* and *Dehalobacterium*, respectively, are highlighted in blue.

The functional potential of the twelve MAGs was assessed based on DRAM annotations. Genes associated with the reductive pentose phosphate cycle, reductive citrate cycle, Wood-Ljungdahl pathway, pentose phosphate pathway, glycolysis, and citrate cycle were detected across all MAGs, whereas none encoded the 3-hydroxypropionate bi-cycle pathway (Figure 3B). Notably, genes involved in the WLP were most abundant in *Dehalobacter* and *D. formicoaceticum*, which also exhibited nearly identical predicted metabolic modules and electron transport chain components, particularly those related to the acetyl-CoA pathway (i.e., CO_2_-to-acetyl-CoA conversion) linked to DCM metabolism. In contrast, the acetyl-CoA pathway was absent from all other MAGs, underscoring the specific potential of *Dehalobacter* and *D. formicoaceticum* to utilize DCM (Figure 3B). In addition, most MAGs encoded F-type ATPase (prokaryotic/chloroplast type), indicating conserved mechanisms for energy transduction. NADH: quinone oxidoreductase, a key component of the bacterial respiratory chain, was detected exclusively in *Dehalobacter* and *D. formicoaceticum* (Figure 3B). Furthermore, genes involved in the conversion of trimethylamine to dimethylamine were identified in three MAGs (UBA5745, *Dehalobacter* and *Dehalobacterium formicoaceticum*), suggesting the potential for anaerobic oxidation of methylated amines (Figure S1).

### Quantitative assessment of active DCM degraders

Total community DNA was extracted from the enrichment consortium at both the onset and completion of DCM degradation, and quantitative PCR (qPCR) targeting the 16S rRNA gene was used to evaluate the growth dynamics of the three putative DCM-degrading populations (*Dehalobacter*, *Dehalobacterium formicoaceticum*, and ‘*Ca*. Dichloromethanomonas elyunquensis’) (Table S2). During both the 3^rd^ and 10^th^ transfers, the copy numbers of the *Dehalobacter* 16S rRNA gene increased substantially. In the 3^rd^ transfer, they rose from (1.1 ± 0.2) x 10^6^ to (8.9 ± 0.2) x 10^7^ copies mL^-1^ (∼81-fold). In the 10^th^ transfer, they increased from (2.1 ± 0.3) x 10^6^ to (2.8 ± 0.41) x 10^8^ copies mL^-1^ (∼136-fold) (Table 1). Notably, *D. formicoaceticum* exhibited detectable but limited growth across all lactate-amended transfers (i.e., transfers 1-10); for example, its abundance increased modestly from (2.6 ± 0.7) x 10^4^ to (3.6 ± 0.3) x 10^5^ copies mL^-1^ in the 3^rd^ transfer and from (1.7 ± 0.5) x 10^4^ to (2.3 ± 0.4) x 10^5^ copies mL^-1^ in the 10^th^ transfer (Table 1). However, throughout the DCM degradation process, the cycle threshold (Ct) values for ‘*Ca.* Dichloromethanomonas elyunquensis’ consistently fell outside the linear detection range of the qPCR assay, indicating that this population was absent from all enrichment cultures. As expected, in lactate-free cultures only *Dehalobacter* proliferated. Its 16S rRNA gene copy numbers increased from (2.5 ± 0.0) x 10^6^ to (2.5 ± 0.4) x 10^8^ copies mL^-1^ in the 11^th^ transfer and from (3.2 ± 0.2) x 10^6^ to (2.1 ± 0.6) x 10^8^ copies mL^-1^ in the 15^th^ transfer over a 7-day incubation period (Table 1). Neither *D. formicoaceticum* nor ‘Ca. Dichloromethanomonas elyunquensis’ was detected under lactate-free conditions, demonstrating that *Dehalobacter* can function as an independent DCM degrader in the absence of auxiliary substrates.

**Table 1.** Growth of *Dehalobacter*, *D. formicoaceticum* and ‘*Ca.* Dichloromethanomonas elyunquensis’ RM with DCM as the growth substrate in different transfer cultures.

| Transfer<sup>a</sup> | *Dehalobacter* (copies) |  | Fold increase<sup>b</sup> | *D. formicoaceticum* (copies) |  | Fold increase | RM (copies)<br>Initial/Final |
| --- | --- | --- | --- | --- | --- | --- | --- |
|  | Initial | Final |  | Initial | Final |  |  |
| 3<sup>rd</sup> | 1.1 ± 0.2 × 10<sup>6</sup> | 8.9 ± 0.2 × 10<sup>7</sup> | 80.9 | 2.6 ± 0.7 × 10<sup>4</sup> | 3.6 ± 0.3 × 10<sup>5</sup> | 13.8 | n.d. |
| 10<sup>th</sup> | 2.1 ± 0.3 × 10<sup>6</sup> | 2.8 ± 0.4 × 10<sup>8</sup> | 133.3 | 1.7 ± 0.5 × 10<sup>4</sup> | 2.3 ± 0.6 × 10<sup>5</sup> | 13.5 | n.d. |
| 11<sup>th</sup> | 2.5 ± 0.0 × 10<sup>6</sup> | 2.5 ± 0.4 × 10<sup>8</sup> | 100.0 | n.d.<sup>c</sup> | n.d. | n.d. | n.d. |
| 15<sup>th</sup> | 3.2 ± 0.2 × 10<sup>6</sup> | 2.1 ± 0.6 × 10<sup>8</sup> | 65.6 | n.d. | n.d. | n.d. | n.d. |

<sup>a</sup> The initial amount of DCM was completely degradation at the end of incubation.
<sup>b</sup> Fold increase in 16S rRNA gene copies is reported as averages of triplicate cultures.
<sup>c</sup> n.d. indicates the target gene was not detected.

### Draft genomes of the DCM-degrading *Dehalobacter* and *Dehalobacterium*

The draft genomes of *Dehalobacter* (hereafter referred to as strain J1) and *D. formicoaceticum* (strain J2) comprised 45 and 104 contigs, respectively, with total genome sizes of 4,311,525 bp (44.5% G+C content, N_50_= 258,283 bp) for strain J1 and 3,269,929 bp (43.2% G+C content, N_50_=70,896 bp) for strain J2 (Table S1). CheckM analysis indicated that strain J1 genome was 99.8% completeness with 1.9% contamination, whereas strain J2 exhibited 91.8% completeness and 1.0% contamination. Genome annotation using RAST predicted 4,201 protein-coding sequences (CDSs) and 52 non-coding RNAs in strain J1, and 3,332 CDSs and 45 non-coding RNAs in strain J2. No plasmids were detected in either genome. Comparative genomic analyses revealed that strain J1 shared moderate similarity with members of the genus *Dehalobacter* (ANI: 72.0-73.7%; AAI: 73.2-75.4%). In contrast, substantially lower relatedness was observed between strain J1 and previously reported anaerobic DCM degraders (strains DMC, RM, EZ94, and DCMF), with ANI and AAI values ranging from 64.6-69.1% and 52.7-68.1%, respectively (Table S3). Phylogenetic analysis based on 16S rRNA gene sequences placed strain J1 within the genus *Dehalobacter*, sharing sequence identities of 94.3%, 96.7%, and 96.8% with *Dehalobacter* sp. TBBPA1, *Dehalobacter* sp. MS, and *Dehalobacter* sp. UNSWDHB, respectively (Figure 4A). In comparison, strain J1 shared <95% 16S rRNA gene sequence identity with members of the genus *Syntrophobotulus* ≤ 93% identity with *Desulfosporosinus* species (Figure 4A). Whole-genome phylogenomic analysis further resolved strain J1 as a distinct lineage most closely related to *Dehalobacter restrictus* DSM 9455 (Figure S3). As expected, strain J2 clustered within an independent 16S rRNA gene branch together with *D. formicoaceticum* strain DMC, sharing 99.9% sequence identity (Figure 4A). Consistently, ANIb values of 98.9% and 98.0% were obtained between strain J2 and the DCM-degrading *D. formicoaceticum* strains EZ94 and DMC, respectively, clearly exceeding the accepted species delineation threshold and confirming its affiliation with *D. formicoaceticum* (Table S3). Based on these genomic and phylogenetic characteristics, strain J1 was proposed as ‘*Candidatus* Dehalobacter formatiformans’ strain J1, representing a novel candidate species within the genus *Dehalobacter*, whereas strain J2 was identified as *D. formicoaceticum* strain J2, a novel strain of the species *D. formicoaceticum*.

**Figure 4.**
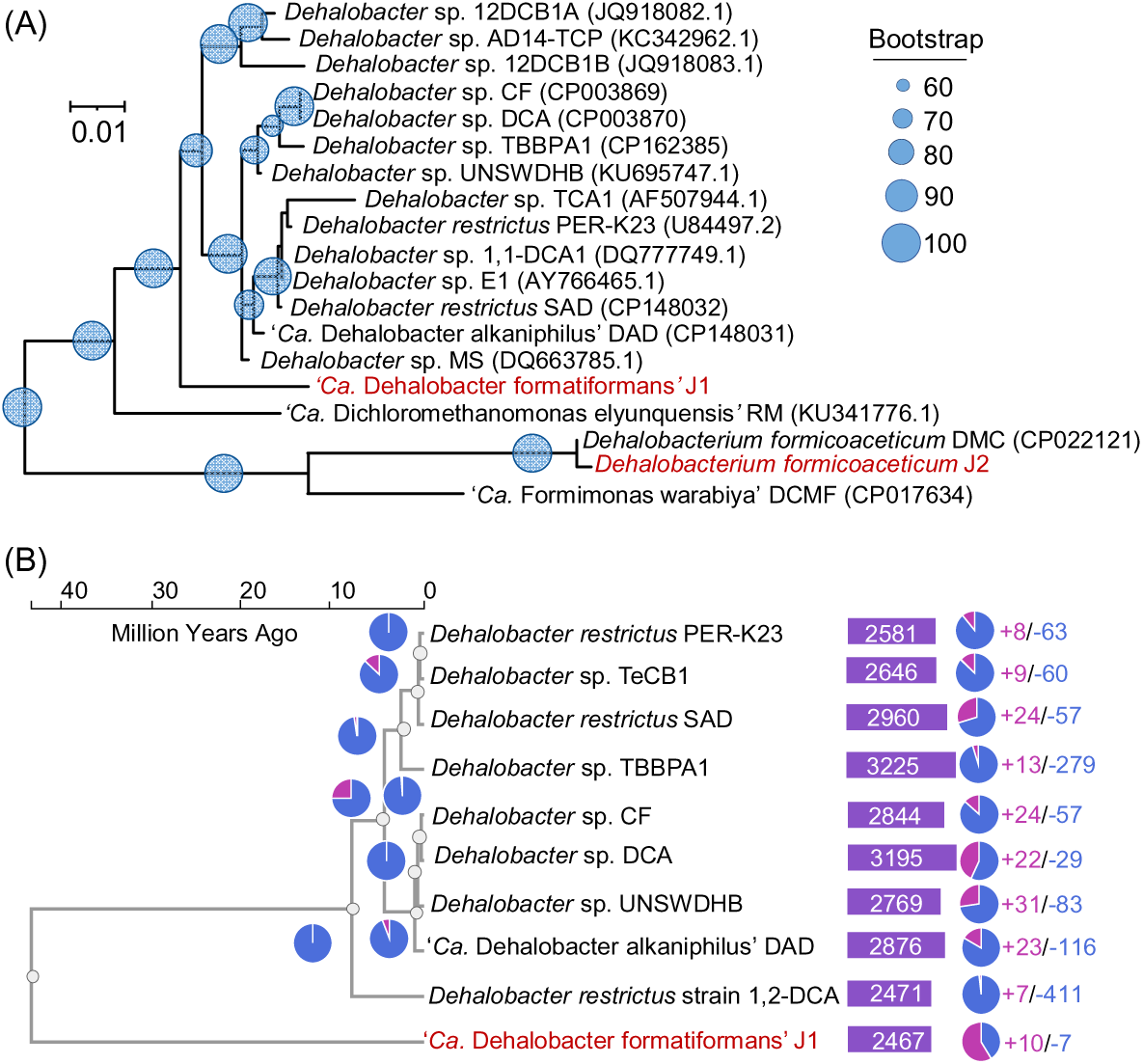
Phylogenetic relationship and evolutionary divergence of strain J1 and related bacteria. (A) Neighbor-joining phylogenetic tree based on 16S rRNA gene sequences showing the relationship among ‘Ca. Dehalobacter formatiformans’ J1, *Dehalobacterium formicoaceticum* J2, representative members of the genus *Dehalobacter*, and previously reported DCM degraders. GenBank accession numbers are indicated in parentheses. Bootstrap values (>60%) based on 1,000 replications are shown at the nodes (filled circles). Scale bar represents 0.01 substitutions per nucleotide base. (B) Gene family variation and species divergence analysis of strain J1 and other *Dehalobacter* species based on concatenated whole-protein sequences. Branch lengths indicate estimated divergence times (MYA) inferred using CAFE5 with orthologous gene clusters identified by OrthoVenn3.

Time Tree analysis based on concatenated whole-protein sequences suggested that the ancestor of ‘*Ca.* Dehalobacter formatiformans’ strain J1 emerged between the Eocene to the late Miocene (approximately 43.33-7.96 million years ago [MYA]), predating the estimated emergence (0.46-7.96 MYA) of obligate organohalide respiring *Dehalobacter* (Figure 4B). Comparative gene family analysis indicated that strain J1 underwent more gene family contractions (10 clusters) than expansions (7 clusters) relative to *Dehalobacter restrictus* strain 1,2-DCA. Conversely, all other *Dehalobacter* genomes exhibited pronounced contraction of 39 gene families compared to strain J1, predominantly involving methanogenesis-related functions (Figure 4B). Around 4.38 MYA, the eight examined *Dehalobacter* genomes diverged into two distinct clades. Clade A comprised *D. restrictus* strains PER-K23 and SAD, *Dehalobacter* sp. TcCB1, and TBBPA1, whereas clade B included *Dehalobacter* sp. CF, DCA, UNSWDHB and ‘Ca. Dehalobacter alkaniphilus’ DAD. Notably, all members of clade B, including obligate OHRB, have been reported to encode partial or complete *mec* cassettes as well as a complete WLP (Figure 4B). Together, these results suggest that the potential for DCM degradation in *Dehalobacter* may have preceded the evolution of obligate organohalide respiration and gradually diminished during subsequent lineage diversification. This metabolic capability could represent an ancestral adaptation to naturally occurring DCM in anoxic, CO_2_-rich environments of the early Earth.

### Genetic characteristics related to anaerobic degradation of DCM

To investigate the genetic basis underlying anaerobic DCM degradation, we compared genome collinearity and gene content of strains J1 and J2, the two microorganisms most likely responsible for DCM catabolism in the consortium, with five experimentally verified anaerobic DCM-degrading strains (i.e., strains RM, DCMF, DMC, DAD and EZ94). Across all examined genomes, a total of 4,366 gene clusters were identified, of which 731 clusters were shared among all strains (Figure 5A). The genomes of ‘*Ca*. Dehalobacter formatiformans’ strain J1, ‘*Ca*. Dehalobacter alkaniphilus’ strain DAD, and ‘Ca. Dichloromethanomonas elyunquensis’ RM shared 80 gene clusters comprising 429 coding sequences, which were primarily associated with positive regulation of transcription, cellular oxidant detoxification, sporulation, and peptidoglycan catabolic processes. In contrast, *D. formicoaceticum* strains J2, EZ94 and DMC shared 717 gene clusters enriched in functions related to serine-type endopeptidase, iron ion homeostasis, plasma membrane components, and DNA restriction-modification system (Figure 5A). Notably, except for strain RM, no reductive dehalogenase (RDase) genes were detected in any of the other anaerobic DCM-degrading genomes, including strains J1 and J2. Interestingly, although strains J1 and J2 each harbored only a partial set of *mec* genes, together they encompass a complete *mec* cassette. Specifically, homologs of *mecA, mecB, mecC, mecH* (two copies), and *mecI* (two copies) were identified in the genome of strain J1, exhibiting amino acid (aa) identities of 84.3-99.8%, 85.9-97.2%, 87.9-99.1%, 43.1-97.2% and 75.6-100%, respectively, relative to corresponding proteins from previously reported DCM degraders. In contrast, strain J2 encoded *mecC, mecD, mecE, mecF, mecG, mecI,* and *mecJ*, with 91.0-98.1%, 81.5-97.8%, 87.5-99.7%, 91.0-99.7%, 77.2-100% and 51.9-96.0% aa identities, respectively. None of the remaining MAGs contained annotated *mec* genes (Figure 5B). Beyond potential limitations imposed by metagenomic assembly, this complementary distribution pattern suggests a potential for conditional or context-dependent metabolic complementation between strains J1 and J2 during DCM degradation. Consistent with other characterized DCM degraders, both strains J1 and J2 encoded a complete set of genes for the WLP, supporting their capacity to metabolize DCM into acetate and formate via the carbonyl and methyl branches of this pathway. In particular, all genes in the WLP of strain J1 and strain J2 were almost identical with those in *Dehalobacter* sp. TBBPA1, and *Dehalobacterium formicoaceticum*, respectively (Table S4).

**Figure 5.**
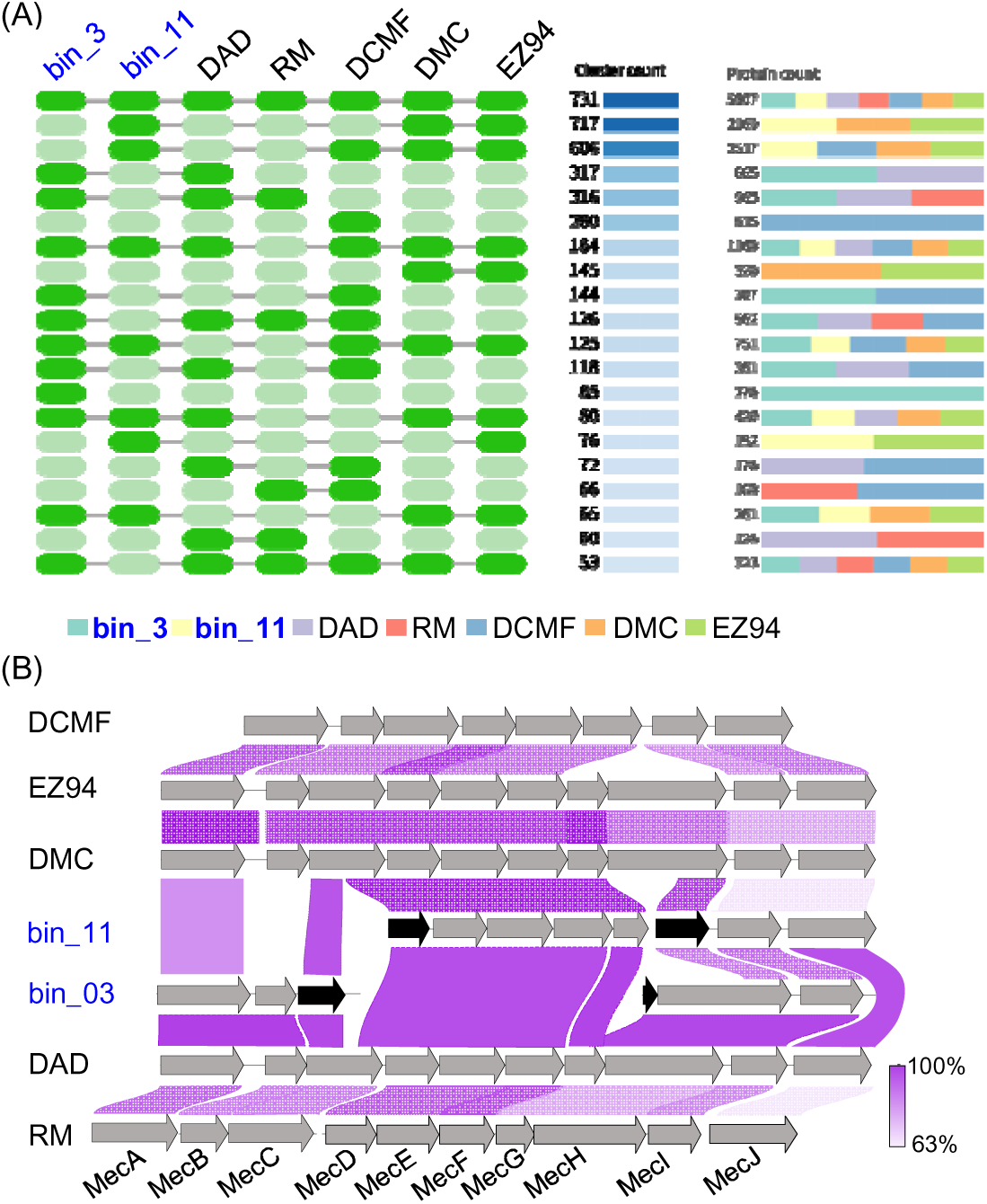
Comparative genomic analysis of DCM-degrading bacteria. (A) Orthologous gene clusters shared among seven DCM-degrading bacteria. Filled and light-green blocks indicate the presence of orthologous gene clusters in each genome. A total of 731 orthologous gene clusters is shared by all seven bacteria. (B) Gene synteny of the *mec* gene cluster among DCM-degrading bacteria. Purple shading indicates the protein sequence identity of MecA-J between species. Genome comparisons were performed using MUMmer. Grey and black arrows represent complete and interrupted Mec proteins, respectively.

Cobamides are essential prosthetic groups for methyltransferases that catalyze methyl-transfer reactions and are therefore thought to play a central role in anaerobic DCM degradation [44]. However, direct evidence for *de novo* cobamide biosynthesis by anaerobic DCM degraders has remained limited. Comparative genomic analysis of currently known anaerobic DCM-degrading bacteria revealed two distinct groups based on their cobamide biosynthetic potential. The first group, represented by strains RM, DAD, and J1, appear to consist of cobamide-auxotrophic organisms. Strain RM lacks more than 70% of the genes required for cobamide biosynthesis, and strains DAD and J1 also lack several key genes (Figure 6A, Table S5). Notably, strain J1 lacks *cysG*, *cbiK* and *cbiZ*, which are responsible for the conversion of uroporphyrinogen III to precorrin-2, cobalt chelation of the corrin ring and cobamide remodeling, respectively [45–47]. In contrast, the second group, including strains DCMF, EZ94, DMC, and J2, retains most genes required for cobamide biosynthesis and appears to have the potential for *de novo* cobamide production (Figure 6A, Table S5). In these organisms, the only consistently missing gene is *cbiK*, while other putative cobalt-chelation genes, such as *cbiX* and *cobN*-like proteins, may perform a similar function [48, 49]. Cultivation experiments in the eighteenth transfer further demonstrated the cobamide dependence of DCM degradation in strain J1. In the absence of exogenous vitamin B_12_, DCM degradation in the consortium completely stalled, with DCM concentrations remaining nearly unchanged over a 60-day incubation period (Figure 6B). In contrast, supplementation with 50 μg/L vitamin B_12_ markedly stimulated DCM removal, resulting in complete degradation within approximately six days at an average rate of 263.7 ± 6.3 μM per day, accompanied by the consumption of approximately 50% of the supplied vitamin B_12_ (Figure 6C). These results clearly indicate that DCM degradation in this strain J1-dominated consortium is strictly cobamide-dependent process.

**Figure 6.**
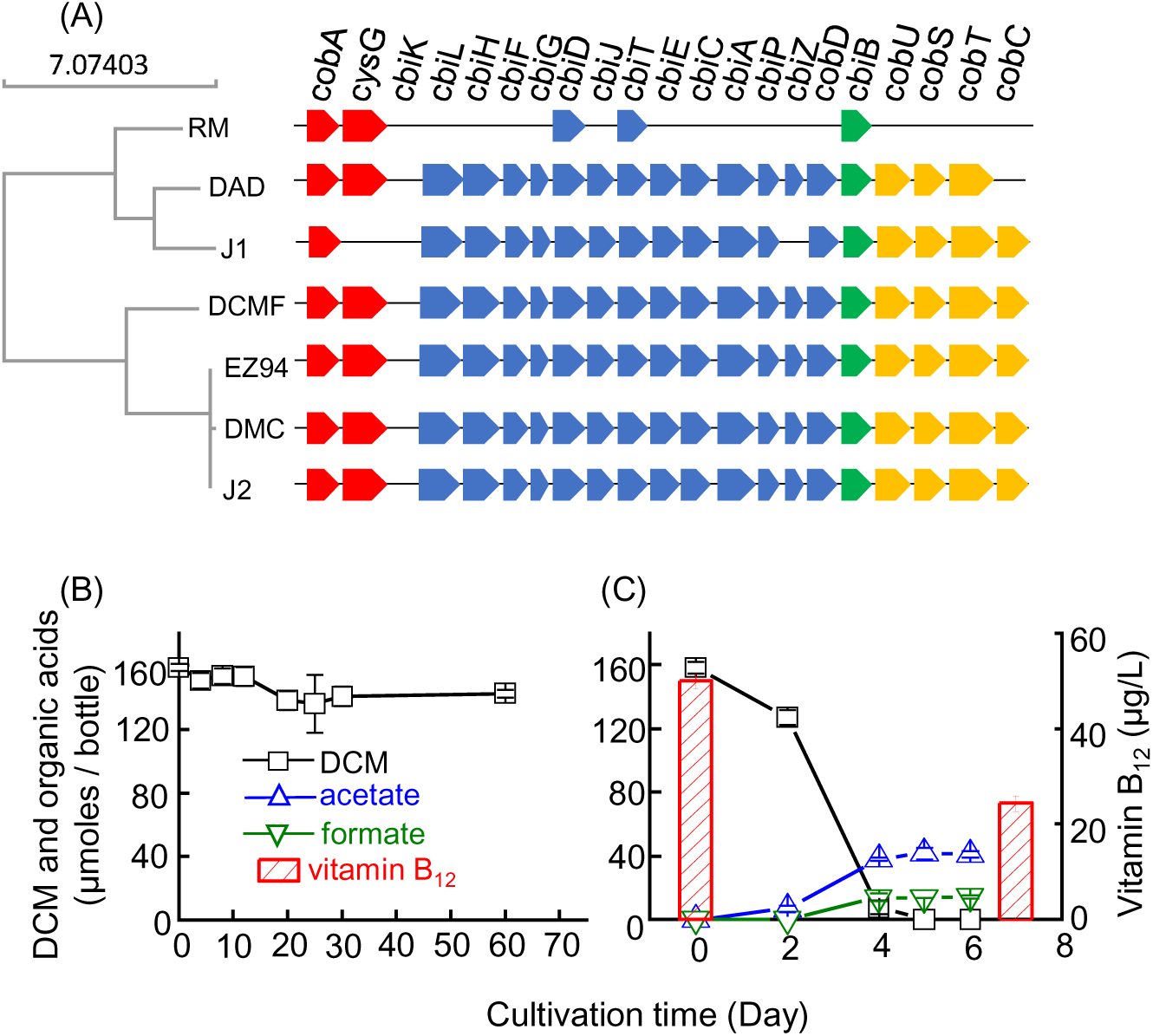
Cobamide biosynthesis potential in DCM-degrading bacteria. (A) Genetic organization of cobamide biosynthesis in seven DCM-degrading bacteria. Genes are represented by colored arrows: red arrows indicate the biosynthesis uroporphyrin III to sirohydrochlorin; blue arrows indicate genes for corrin ring biosynthesis; green arrows indicate cobamide linker biosynthesis genes; yellow arrows indicate genes involved in nucleotide loop assembly. Regions lacking arrows indicate the absence of the corresponding genes. (B) DCM degradation in the transfer culture maintained without exogenous vitamin B_12_ supplementation. (B) DCM degradation and vitamin B_12_ consumption in the corresponding transfer culture supplemented with 50 μg/L vitamin B_12_ during the eighteenth transfer. Error bars represent the standard deviations of three biological replicates.

### Description of ‘Candidatus Dehalobacter formatiformans’ sp. nov

‘Candidatus Dehalobacter formatiformans’ (for.ma.ti.for’mans. L. n. formate (formic acid salt); L. part. adj. formans, forming or producing; N.L. part. adj. formatiformans, formate-forming. The species epithet refers to the bacterium’s ability to produce formate and acetate during the anaerobic degradation of DCM. ‘Candidatus Dehalobacter formatiformans’ represents a strictly anaerobic bacterium belonging to the genus *Dehalobacter*. The organism in a mixed culture derived from river sediment collected from the Xi River, Shenyang, China (41.6628°N, 123.1055°E), utilizes DCM as its sole organic carbon and energy source, converting it into formate and acetate via a cobamide-dependent pathway. Growth occurs at 20-30 °C and circumneutral pH in bicarbonate-buffered medium under anoxic conditions, with DCM degradation ceasing in the absence of exogenous cobamide. Phylogenetic, genotypic and phenotypic characteristics place strain J1 in the *Dehalobacter* genus within the DCM-degrading Firmicutes, and we propose that strain J1 represents a new species, ‘Candidatus Dehalobacter formatiformans’.

## Discussion

The results of this study expand the current understanding of the ecological and metabolic roles of *Dehalobacter* in anaerobic environments. Members of this genus have long been regarded primarily as obligate OHRB specialized in reductive dechlorination of multi-carbon organohalogens. However, the findings presented here suggest that *Dehalobacter* may also participate in the transformation of chlorinated C1 compounds through metabolic strategies fundamentally different from classical organohalide respiration. In contrast to reductive dehalogenation processes that couple halogen removal to respiratory energy conservation, the pathway observed in this system appears to integrate DCM-derived C1 units into central carbon metabolism via methyl-transfer reactions and the WLP, resulting in the formation of acetate and formate. This metabolic configuration indicates that DCM can function not only as a halogenated electron acceptor analogue but also as a fermentable C1 substrate within certain anaerobic bacterial lineages. Such versatility suggests that the capacity for DCM transformation in *Dehalobacter* may represent a previously underappreciated metabolic dimension of this genus and highlights a broader role for C1 halogenated compounds in shaping anaerobic carbon fluxes in subsurface ecosystems.

In line with these considerations, the total carbon fixed from DCM into products and biomass was evaluated based on the 15^th^ lactate-free transfer cultures, where ‘Ca. Dehalobacter formatiformans’ dominated the community. The total fixed carbon (C_fixed) was calculated as the sum of carbon in metabolites and biomass, where metabolite carbon is the carbon in acetate and formate (98.6 μmol). Biomass carbon was calculated from the increase in cell abundance according to the equation 1 [50, 51], where 12.011 is the atomic weight of carbon. The cell carbon content (20 fg C per cell) was estimated based on previous studies of bacterial biomass. The resulting total fixed carbon was 132.0 μmol, giving a carbon fixation efficiency of 84.6% (equation 2). This high efficiency indicates that the majority of carbon driven by anaerobic DCM degradation was incorporated into metabolites and biomass rather than being lost as CO_2_, and it is noteworthy that this anaerobic DCM degradation occurred in the presence of CO_2_, which likely acted as a co-substrate or electron sink, thereby directly contributing to net carbon fixation in the system. By integrating hazardous chlorinated solvents into biomass and metabolite pools, these systems exemplify a circular carbon economy approach, turning waste into valuable products.

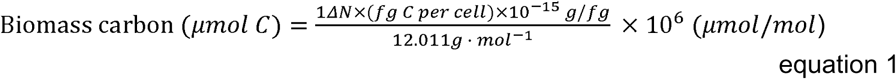

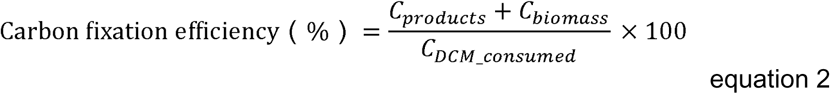

Comparative genomic analysis revealed that all available *Dehalobacter* genomes, including the DCM-degrading strain J1 in this study, possess a complete set of genes encoding the Wood-Ljungdahl pathway, although the functional role of this pathway in most non-DCM-degrading *Dehalobacter* strains remains unclear, and no known *Dehalobacter* isolate was described to grow autotrophically with H_2_ and CO_2_. It has been hypothesized that the WLP might have another role than CO_2_-fixation in *Dehalobacter* strains because some *Dehalobacter* species (e.g., strain TecB1) were isolated and grown in a bicarbonate-free medium [52]. Besides, both chloroform (CF)-respiring and DCM-mineralizing *Dehalobacter* strains share notable genetic commonalities [10, 38, 43]. For instance, although *Dehalobacter* sp. strains UNSWDHB and CF are not capable of DCM mineralization, they still encode part or even a complete *mec* cassette, and their *mec* gene orthologs (particularly *mecA*, *mecB*, and *mecC*) exhibit sequence identities of 93.8-97.4% with genes found in the genomes of DCM-degraders. Conversely, strains like RM and DAD, which do not perform reductive dechlorination, harbor multiple putative RDase genes. These observations suggest a potential functional link between the CF dehalogenations and DCM mineralization, but also an evolutionary divergence within *Dehalobacter*, with different strains adapting to distinct organohalide or C_1_ compound metabolisms.

Cultivation and environmental studies have revealed an increasing number of halogenated compounds that can be used by *Dehalobacter* sp. as electron acceptors including tetrachloroethene, CF, chlorinated ethanes, 2,4,6-trichlorophenol, 2,4,6-tribromophenol, chlorinated benzenes, β-hexachlorocyclohexane (β-HCH, known as lindane) and phthalide [53]. These compounds can be categorized according to their natural origins: C_1_ halogenated compounds such as CM, bromomethane, and DCM are widely produced in nature by fungi, algae, higher plants, and biomass burning, whereas higher halogenated aromatics and alkanes are largely of anthropogenic origin [3]. The natural abundance of C_1_ organohalogens suggests that early microorganisms on Earth could only exploit these compounds as electron acceptors, making it reasonable that the evolution of anaerobic DCM degradation predates that of organohalide respiration (Figure 4B). Moreover, the functionality of DCM degradation may have originated from ancient methanogenic archaea, as evidenced by the presence of an 8,191 bp methanogen-derived integrative fragment (98.9% sequence identity to the *Methanobacterium* sp. MB1 [54] detected in strain J1 genome. The segment encodes five genes, each annotated as peptidoglycan DD-metalloendopeptidase family protein, SNF2 domain protein, cadmium resistance protein CadD, DUF3883 domain-containing protein, site-specific recombinase, DNA invertase Pin and Cadmium-translocating P-type ATPase. The transfer of these genes does not appear to be associated with energy metabolism of DCM, but contributes to the adaptation of microorganisms to the external environment, e.g., resistance to toxic environments with high cadmium, and thus plays a key role in the plasticity and evolutionary evolution of the host genome.

The observed DCM catabolism in this study likely depends on metabolic cooperation not only between J1 and J2 but also on cobamide availability from the surrounding community. Such syntrophic interactions highlight the importance of community context in anaerobic halogenated C_1_ metabolism. These interactions may influence both substrate turnover and the efficiency of carbon incorporation, implying that single-species studies might underestimate the functional potential and ecological significance of complex microbial consortia in sedimentary carbon and halogen cycles. In our enrichment consortium, several cobamide-producing microorganisms (e.g., *Sporomusa* and *Clostridium*) [55] were detected, indicating a potential internal source of cobamide. However, external supplementation of vitamin B₁₂ remained essential for sustaining DCM degradation activity. This phenomenon likely reflects two possibilities: first, cobamide-synthesizing populations occur at low abundance, and thus produce insufficient cobamide to support the community’s overall metabolic demand; second, the cobamide variants synthesized by these organisms may not match the structural requirements of DCM-degrading enzymes. The latter scenario is plausible, as previous studies have demonstrated that many cobamide-dependent metabolisms exhibit strong selectivity toward specific lower-base structures [56]. Elucidating these intricate cobamide exchange and compatibility networks remains a challenging but crucial task for future research.

## Materials and Methods

### Chemicals

DCM (purity ≥99.8%) and chloroform (CF, ≥99.9%) were procured from China National Pharmaceutical Group Co., Ltd. (Shanghai, China). CM (99.5%) and CH_4_ (99.99%) were obtained from Dalian Special Gasses Co., Ltd. (Dalian, Liaoning, China). Vitamin B_12_ (≥97.0%) was procured from Sigma-Aldrich (St. Louis, MO, USA). All other chemicals used in this study were of analytical reagent grade or higher quality.

### Microcosms and enrichment cultures

Microcosms were established as described previously [57] in 160-mL (nominal capacity) serum bottles containing 100 mL of reduced, bicarbonate-buffered mineral salts medium (30 mM, pH 7.3) amended with lactate (5 mM), 2-bromoethanesulfonate (BES, 2 mM) and 0.1 mL of appointed Wolin vitamins containing 50 μg/L vitamin B_12_ [58], and 10 µL neat DCM (*ca.* 156 µmoles/bottle). Approximately 2 g of anoxic sediment material collected from the outlet area of a wastewater treatment plant in the Xi River (41.6628°N, 123.1055°E) [59, 60] as inoculum was transferred into each bottle in an anaerobic glove chamber (95% N_2_-5% H_2_) (Coy Laboratory, Ann Arbor, MI). After ten consecutive transfers under the same incubation conditions as the microcosm, lactate and BES were removed from the subsequent transfer cultures. All vessels were statically incubated at 30 °C in the dark.

### DNA extraction and 16S rRNA gene amplicon sequencing

Culture suspension samples (5 mL) were collected and filtered through 0.22 μm filtration membranes (Millipore, Burlington, MA, USA). Genomic DNA extraction was performed using the HiPure Soil DNA Kit (Magen Biotech, Inc) according to the provided manual. PCR procedures were carried out using the universal primer set V3-V4-F and V3-V4-R with approximately 20 ng of DNA as the template [61]. The resulting amplified products, which contained indexed adapters at the ends, were purified. Subsequently, 2×300 paired-end sequencing with dual reads was performed using the Illumina Miseq PE250/300 Platform (Illumina, San Diego, USA) at Azenta Life Sciences Inc. (South Plainfield, NJ, USA). To process the raw sequences, trimming and pairing were performed using QIIME2-v.2018.11 [62]. The optimized sequences were then subjected to clustering into operational taxonomic units (OTUs) using SILVAngs v1.9.8 with a similarity threshold set at 98% [63].

### Metagenome sequencing, assembly, and binning

Metagenomic sequencing of the 10^th^ DCM-degrading enrichment culture was conducted using the NovaSeq 6000 PE150 sequencer at Novogene Bioinformatics Technology (Beijing, China). Genomic DNA was extracted from the stationary phase cell pellets using the sodium dodecyl sulfate (SDS) method as described [64]. The integrity and size of the extracted DNA were assessed through 1.0% agarose gel electrophoresis at 120 V, while the DNA concentration was measured using a Qubit 2.0 Fluorometer (Thermo Fisher Scientific, Waltham, MA, USA). For library preparation, randomly sheared DNA fragments were generated using the NEBNext® Ultra™ DNA Library Prep Kit for Illumina (NEB, USA). The raw sequencing data underwent initial processing to remove low-quality reads utilizing Trimmomatic v0.38 with default parameters [65]. High-quality reads were then assembled using the JGI Metagenome Assembly Pipeline (https://github.com/kbaseapps/jgi_mg_assembly) [66]. Kaiju v1.7.3 was employed to perform read taxonomic classification [67]. Assembly contigs were binned into species-level metagenome-assembled genomes (MAGs) using Maxbin2-v2.2.4 [68]. Quality assessment and taxonomic assignment of the MAGs were determined using CheckM v1.0.18 [69] and GTDB-Tk classify [70], respectively. MAGs meeting the criteria of ≥ 70% completeness and < 10% contamination, as previously defined [71], were retained. Gene and protein sequences within each MAG were predicted and annotated using DRAM [72]. Whole genome pairwise comparison and annotation of orthologous clusters of DCM degrading strains were performed using OrhoVenn3 [73]. Average nucleotide identity (ANI) and Average amino acid identity (AAI) were calculated using JSpeciesWS [74] and an online calculator form Kostas lab [75], respectively.

### Phylogenetic analysis

To assess the phylogenetic relatedness of strain J1, strain J2, *Dehalobacter* spp, *Dehalobacterium* and other anaerobic DCM degraders. 16S rRNA gene sequences were obtained from the NCBI database based on the BLAST results of both strains. Only sequences showing > 93% similarity to either strain J1 or J2 were selected for phylogenetic tree construction. Multiple sequence alignment of the 16S rRNA gene sequences was performed using Muscle version 3.8 [76]. The aligned sequences were then used to construct a phylogenetic tree using the neighbor-joining method with 1,000 bootstrap replications in Geneious Prime v11.0.

### Quantitative polymerase chain reaction (qPCR)

Organism-specific qPCR assays each targeting the 16S rRNA gene of *Dehalobacter*, *Dehalobacterium formicoaceticum*, and ‘*Ca*. Dichloromethanomonas elyunquensis’, respectively, were used to monitor the growth of the respective populations in the DCM-degrading mixed cultures. DNA was extracted using the TIANamp Soil DNA kit (Tiangen Biotech Inc.) following the manufacturer’s protocol. qPCR primers and probes are listed in Table S2. qPCR analysis followed published protocols and was conducted using an QuantStudio^TM^ 3 Real-Time PCR system (Applied Biosystems, Waltham, MA, USA). Briefly, every 25 µL reaction mixture contained 12.5 μL of 2 × Premix Ex Taq^TM^ master mix (Takara Bio Inc.), 2 μL of DNA template, forward and reverse primers and probe at final concentrations of 200 nM each, and 0.25 μL of 50 × ROX reference Dye II (Takara Bio Inc.). The PCR thermal cycling protocol was as follows: 50°C for 2 min, then at 95°C for 10 min, followed by 40 cycles of denaturation at 95°C for 15 sec and annealing and extension at 60°C for 1 min. Calibration curves used triplicate serial 10-fold dilutions of plasmid DNA (pUC-SP vector, Sangon Biotech Inc.) carrying three fragments of the 16S rRNA genes of ‘*Ca*. Dehalobacter formatiformans’ strain J1, *Dehalobacterium formicoaceticum*, and ‘*Ca*. Dichloromethanomonas elyunquensis’, and spanned the concentration range from 6.24 x 10^9^ to 6.24 x 10^2^, 2.99 x 10^9^ to 2.99 x 10^2^, and 1.27 x 10^9^ to 1.27 x 10^3^ target gene copies per assay tube, respectively (Figure S1).

### Sequence data deposition

Metagenome sequencing raw reads were deposited in the Sequence Read Archive (SRA) under accession number SRR37122118. The BioSample and BioProject accession numbers are SAMN55067445 and <u>PRJNA1418947</u> for strain J1, SAMN55067543 and <u>PRJNA1418954</u> for strain J2, respectively.

## Acknowledgements

This work was supported by the Intergovernmental International Cooperation in Science and Technology Innovation of National Key Research and Development Project of China (Grant No. 2023YFE0122000), the National Natural Science Foundation of China (42407189, 42307186), the Open Fund Project of the State Environmental Protection Key Laboratory of Soil Environmental Management and Pollution Control (MEESEPC202508), the Shanghai Frontiers Science Center of Polar Science (Grant No. SOO2025-01), the Open Funding of State Key Laboratory of Coal Mine Disaster Prevention and Control (Grant No. 2024SKLKF03), the International Science and Technology Cooperation Project of Department of Science and Technology of Liaoning Province (Grant No. 2024JH2/101900003).

## Author Contributions

Y. Y. and H.J. conceived the study and designed the research. H.J. performed the enrichment cultivation, physiological experiments, and molecular analyses. H.J., X.L., X.W., H.W., J.W., K.S., G.L., T.Z., and S.H. contributed to cultivation experiments, chemical analyses, and data acquisition. H.J. conducted metagenomic assembly, genome analysis, and data interpretation. All authors contributed to discussion and interpretation of the results. H.J. drafted the manuscript. Y.Y. and J.Y. revised the manuscript. M.M. and F.E.L. provided critical feedback and substantially improved the manuscript.

