## Supplementary material for "Cobamide-Dependent Dichloromethane Fermentation by *Dehalobacter* Reveals a Hidden Acetogenic Route for Organohalide Biotransformation": Table S1

^1^ Institute of Applied Ecology, Chinese Academy of Sciences; Shenyang, Liaoning 110016, China; ^2^ Department of Civil, Environmental and Construction Engineering, University of Hawaii at Manoa, Honolulu, Hawaii 96822, United States; .^3^ Key Laboratory of Forest Ecology and Silviculture, Institute of Applied Ecology, Chinese Academy of Sciences, Shenyang, Liaoning, 110016, China. ^4^ University of Chinese Academy of Sciences, Beijing, China, 100049; ^5^ School of Biotechnology and Biomolecular Sciences, University of New South Wales, Sydney, 2052, Australia. ^6^ Department of Civil and Environmental Engineering, University of Tennessee, Knoxville, Tennessee 37996, USA; ^7^ Department of Biosystems Engineering and Soil Science, University of Tennessee, Knoxville, Tennessee 37996, USA; ^8^ Department of Biochemistry & Cellular and Molecular Biology, University of Tennessee, Knoxville, Tennessee 37996, USA.

*** Corresponding authors**

Yi Yang, Institute of Applied Ecology, Chinese Academy of Sciences, 512 South Building, 72 Wenhua Road, Shenyang, Liaoning 110016, China, Phone: +86-2483970426,

**ORCID**

Yi Yang: 0000-0002-3519-5472

Huijuan Jin: 0000-0003-0686-2309

Xiuying Li: 0000-0003-3555-7418

Michael Manefield: 0000-0002-1880-0888

Frank E. Löffler: 0000-0002-9797-4279

Jun Yan: 0000-0001-6883-8529

**Supplementary Information Text**

**Identification of *mec* genes in strains J1 and J2.** The *mec* gene cluster from the *Dehalobacterium* *formicoaceticum* strain DMC genome was employed as a reference for comparative genomic alignment with strains J1 and J2 to identify their respective *mec* gene counterparts in the Geneious prime v11.0. The synteny plot was generated using the gbk files as input for pyGenomeViz v1.6.1 (mode: MUMmer; identity threshold of 50%) (1).

**Vitamin B_12_	concentration analysis.** Vitamin B_12_ in culture supernatants was extracted as described (2). Briefly, 1 mL of culture was centrifuged at 20,000 x *g* for 3 min and the supernatant was filtered through a 0.22-μm polyethersulfone syringe filter. A 0.5 mL aliquot of the filtrate was adjusted to pH ~ 5.5 by adding 1.75 μL glacial acetic acid, followed by reaction with 20 mM KCN under ambient air in sealed tubes at 100 ^o^C for 60 min. After cooling, samples were centrifuged again (20,000 × g, 10 min), and 400 μL of supernatant was collected for LC-MS/MS within 2 h to minimize interference from reducing agents (e.g., Na_2_S, DTT). LC-MS/MS was performed using a Vanquish^TM^ UHPLC coupled to a TSQ Quantis^TM^ mass spectrometer (Thermo Fisher Scientific, USA). A 10 μL aliquot was injected onto a Hypersil GOLD™ C18 column (2.6 μm, 2.1 × 100 mm) at 30 °C and 0.3 mL min⁻¹. Eluents were (A) 0.1% formic acid in water and (B) acetonitrile. The gradient was 95% A → 85% A (2.8 min) → 75% A (1.7 min) → 30%A (1.5 min), followed by re-equilibration to 95% A for 1.5 min. The mass spectrometer operated in positive ESI mode with a spray voltage of 3.8 kV, capillary temperature of 350 ^o^C, sheath gas 40, and auxiliary gas 10 (arbitrary units). Vitamin B_12_ was quantified in SRM mode using transitions from the doubly charged precursor ion to product ions representing the lower base, phosphoriboside, and free base fragments. Data were processed with Xcalibur™ 3.0 (Thermo Fisher Scientific).

**P****hylogenetic analysis.** The species divergence time tree and the expansion and contraction of gene families based on protein sequences from 10 selected *Dehalobacter* spp. species were analyzed by OrthoVenn3 with CAFE5 (3). The whole proteome (amino acid sequences)-based phylogenetic tree was obtained from the Type Strain Genome Server (<https://tygs.dsmz.de>) with a distance algorithm of greedy-with-trimming (4). Species tree analysis of DCM anaerobic degrading bacteria was conducted by OrthoVenn3 with method of maximum likelihood and evolution model of JTT+CAT.

**Analytical methods**. Chlorinated solvents were quantified using an Agilent 7890A gas chromatograph (GC) equipped with a flame ionization detector and an Agilent DB-624 column (60 m × 0.32 mm × 1.8 μm), as described in the reference (5). The GC analysis allowed for the measurement of chlorinated solvents. Average degradation rate was calculated based on the consumption of DCM during the entire degradation process. Organic acids were analyzed using an Agilent 1260 HPLC system. An Aminex HPX-87H column from Bio-Rad (Hercules, CA) was used with 4 mM H_2_SO_4_ as the eluent at a flow rate of 0.6 mL/min. The quantification of organic acids was performed using a diode array detector set to 210 nm (6).

**Figures and Tables**

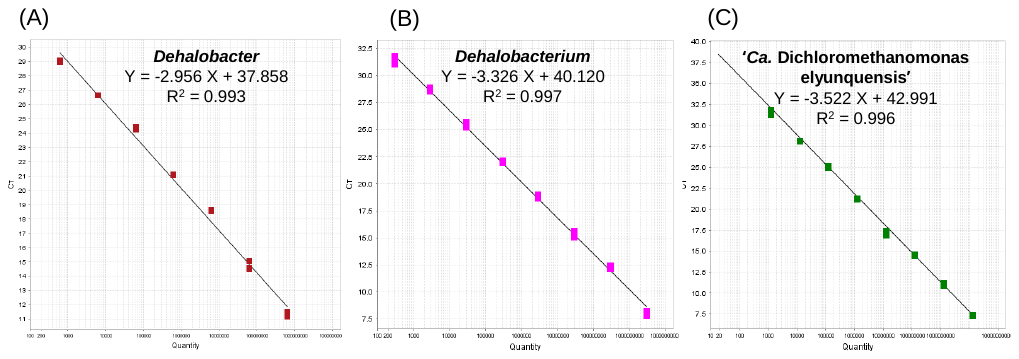

**Figure S1.** **The standard curve of the qPCR assay targeting the 16S rRNA gene of three DCM-degrading bacteria.** (A) *Dehalobacter*; (B) *Dehalobacterium*; (C) ‘*Ca*. Dichloromethanomonas elyunquensis’.

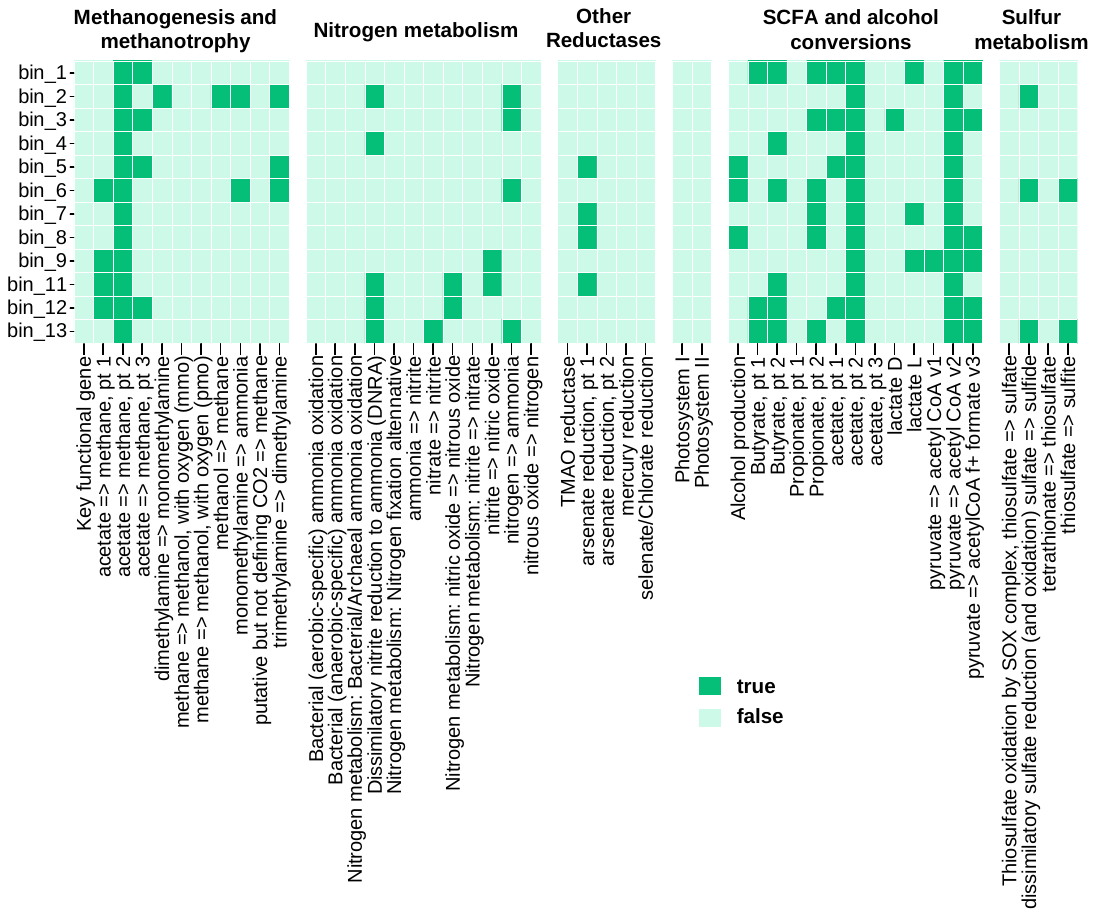

Figure S2. **Metabolic potential of qualified MAGs recovered from the DCM-degrading enrichment cultures.** Presence/absence of 1-2 key genes required for each metabolic process was determined based on DRAM annotation profiles.

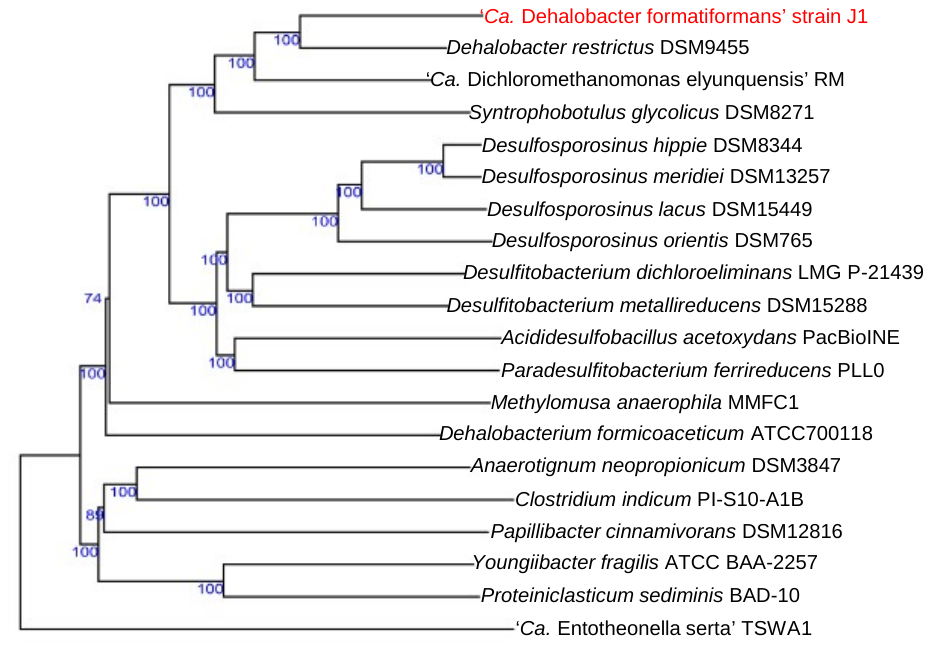

**Figure S3.** Genome-based phylogenetic tree reconstructed using FastME 2.1.6.1 based on Genome BLAST Distance Phylogeny (GBDP) distances derived from whole-genome sequences. Branch lengths correspond to GBDP distance formula d5. Values at the nodes represent GBDP pseudo-bootstrap support (>60%) based on 100 replicates. Genome-based taxonomic analysis was conducted using the Type (Strain) Genome Server (TYGS) (https://tygs.dsmz.de).

**Table S1. Bin summaries from metagenomic binning.**

| **UniteM Bin** | **Classification** | **Completeness**  **(%)** | **Contamination**  **(%)** | **Genome Size (bp)** | **No. Contigs** | | **N50** | **L50** |
| --- | --- | --- | --- | --- | --- | --- | --- | --- |
| bin_01 | f_Clostridiaceae | 96.90 | 3.61 | 3,140,965 | 83 | 69,977 | | 15 |
| bin_02 | f_UBA5745 | 89.14 | 1.88 | 2,531,520 | 44 | 96,829 | | 10 |
| bin_03 | g_*Dehalobacter* | 99.81 | 1.86 | 4,311,525 | 45 | 258,283 | | 7 |
| bin_04 | F_Oscillospiraceae | 94.21 | 0.67 | 2,004,529 | 154 | 16,319 | | 33 |
| bin_05 | s_*Anaerotignum propionicum* | 81.18 | 2.35 | 2,733,338 | 365 | 9,141 | | 87 |
| bin_06 | g_*Propionicimonas* | 83.38 | 6.47 | 3,401,430 | 555 | 7,184 | | 147 |
| bin_07 | g_*Lentimicrobium* | 97.27 | 4.03 | 4,735,443 | 187 | 66,124 | | 23 |
| bin_08 | g_*Petrimonas* | 90.98 | 6.08 | 2,717,830 | 406 | 8,214 | | 104 |
| bin_09 | c_Negativicutes | 86.01 | 2.90 | 3,363,904 | 427 | 10,290 | | 92 |
| bin_11 | s_*Dehalobacterium formicoaceticum* | 91.78 | 1.02 | 3,269,929 | 104 | 70,896 | | 15 |
| bin_12 | f_Anaerotignaceae | 89.04 | 9.35 | 3,208,915 | 628 | 5,486 | | 184 |
| bin_13 | g_Bact-08 | 95.37 | 5.75 | 2,632,673 | 122 | 52,182 | | 18 |

**Table S2. The list of primers or probes used in this study.**

| **Primer** | **Sequence (5’-3’)** | **Target gene** | **Reference** | |
| --- | --- | --- | --- | --- |
| 27F | AGAGTTTGATCCTGGCTCAG | Bacterial 16S rRNA | | (7) |
| 1492R | GGTTACCTTGTTACGACTT |  |  |  |
| V3-V4-F | CCTACGGRRBGCASCAGKVRVGAAT ^a^ | Bacterial 16S rRNA | | (8) |
| V3-V4-R | GGACTACNVGGGTWTCTAATCC ^a^ |  |  |  |
| Dhb1200F | CCTTAAGAGATTAGGGAGTGCC | *Dehalobacter* (bin_03)  16S rRNA | | This study |
| Dhb1340R | CTCAGCTTTACCTGTTAGCAAC |  |  |  |
| Dhb1236P | 6FAM-GACGCAGGTGGTGCATGGTTGTCG-MGB ^b,c^ |  |  |  |
| Defo1022F | CCTTGACAGTCATGGAAACATG | *Dehalobacterium*  *formicoaceticum* 16S rRNA | | This study |
| Defo1174R | GCCCACCTTATATGCTGGC |  |  |  |
| Defoq1070P | 6FAM-GACGCAGGTGGTGCATGGTTGTCG-MGB ^b,c^ |  |  |  |
| Diel1019F | GTCTGAGGAGACTCGGATG | *‘Ca.* Dichloromethanomonas elyunquensis’  16S rRNA | | This study |
| Diel1158R | CACCTTTACGTGCTGGTAACTG |  |  |  |
| Diel1056P | 6FAM-GACGCAGGTGGTGCATGGTTGTCG-MGB ^b,c^ |  |  |  |

^a^ Degenerate bases, R=A/G, B=C/G/T, S=C/G, K=G/T, V=A/C/G, N=A/C/G/T, W=A/T.

^b^ 6FAM, 6-carboxyfluorescein; MGB, minor groove binder moiety.

^c^ The probes of Dhb1236P, Defoq1070P and Diel1056P have the same sequence.

**Table S3**. The average nucleotide identity (ANI) values (%) (lower left half) and the average amino acid identity (AAI) (%) (upper right half) of strains J1 and J2 with other strains in the genus of *Dehalobacter* and five DCM dergarders from previous studies (9-12). Strains: 1. *S*train J1; 2. *Dehalobacter restrictus* strain 12DCA (CP046996); 3. *Dehalobacter* sp. UNSWDHB (AUUR01000000); 4. ‘*Candidatus* Dehalobacter alkaniphilus’ DAD (CP148031); 5. *Dehalobacter* sp. 14DCB1 (PNXX01000000); 6. *Dehalobacter* sp. CF (CP003870); 7. *Dehalobacter* sp. TeCB1 (MCHF01000000); 8. *Dehalobacter* *restrictus* SAD (CP148032); 9. *Dehalobacter* sp. TBBPA1 (CP162385); 10. *Dehalobacterium formicoaceticum* strain DMC (CP022121); 11. *D. formicoaceticum* EZ94 (JANPWE010000000); 12. ‘*Candidatus* Dichloromethanomonas elyunquensis’ (LNDB01000000); 13. ‘*Candidatus* Formimonas warabiya’ DCMF (CP017634); 14. Strain J2.

| **Strains** | **G+C（%）** | **The average nucleotide identity (ANI) and average amino acid identity (AAI) (%)** | | | | | | | | | | | | | |
| --- | --- | --- | --- | --- | --- | --- | --- | --- | --- | --- | --- | --- | --- | --- | --- |
|  |  | 1 | 2 | 3 | 4 | 5 | 6 | 7 | 8 | 9 | 10 | 11 | 12 | 13 | 14 |
| 1 | 44.5 | * | 73.2 | 74.4 | 75.4 | 73.3 | 73.8 | 73.8 | 74.8 | 73.7 | 54.1 | 53.9 | 68.1 | 52.9 | 52.7 |
| 2 | 44.5 | 72.8 | * | 95.0 | 95.0 | 92.3 | 94.5 | 98.5 | 98.4 | 96.5 | 54.6 | 54.7 | 70.7 | 53.0 | 53.9 |
| 3 | 44.9 | 72.5 | 94.3 | * | 99.0 | 92.6 | 98.9 | 95.5 | 95.4 | 94.9 | 55.3 | 55.3 | 70.2 | 53.2 | 54.6 |
| 4 | 44.9 | 73.3 | 94.1 | 98.3 | * | 92.1 | 99.1 | 95.3 | 95.4 | 95.2 | 55.1 | 54.7 | 70.6 | 53.3 | 54.1 |
| 5 | 44.0 | 73.7 | 90.9 | 91.1 | 90.6 | * | 93.0 | 95.3 | 91.6 | 93.3 | 55.9 | 55.7 | 70.1 | 53.2 | 55.2 |
| 6 | 44.3 | 71.9 | 94.4 | 98.8 | 98.8 | 91.7 | * | 95.0 | 95.4 | 95.2 | 56.2 | 55.5 | 70.3 | 53.4 | 54.6 |
| 7 | 44.0 | 72.4 | 97.8 | 94.5 | 94.5 | 91.1 | 94.1 | * | 98.6 | 97.1 | 53.9 | 53.7 | 70.7 | 53.3 | 53.3 |
| 8 | 44.2 | 73.1 | 98.2 | 94.7 | 94.9 | 90.7 | 94.7 | 98.5 | * | 96.7 | 54.4 | 54.5 | 70.9 | 53.3 | 54.0 |
| 9 | 44.6 | 72.3 | 95.9 | 93.7 | 93.7 | 91.6 | 93.7 | 96.3 | 96.2 | * | 53.8 | 53.9 | 70.8 | 53.1 | 53.7 |
| 10 | 43.2 | 65.7 | 67.3 | 67.8 | 67.1 | 67.9 | 68.5 | 66.2 | 66.7 | 65.9 | * | 98.9 | 53.5 | 66.8 | 99.1 |
| 11 | 43.2 | 65.1 | 66.7 | 66.8 | 66.0 | 67.6 | 67.3 | 65.3 | 66.0 | 65.3 | 98.7 | * | 53.9 | 67.1 | 99.2 |
| 12 | 43.0 | 69.1 | 70.9 | 70.8 | 70.9 | 70.2 | 70.5 | 70.7 | 71.0 | 70.8 | 65.2 | 65.2 | * | 53.3 | 53.1 |
| 13 | 46.4 | 64.8 | 64.9 | 65.1 | 65.4 | 65.3 | 65.3 | 65.3 | 65.4 | 65.0 | 69.5 | 69.2 | 64.2 | * | 67.0 |
| 14 | 43.2 | 64.6 | 66.2 | 66.6 | 66.3 | 67.4 | 66.4 | 65.7 | 66.4 | 65.4 | 98.0 | 98.9 | 65.0 | 69.0 | * |

**Table S4.** Wood-Ljungdahl pathway genes identified in in the genomes of 'Ca. Dehalobacter formatiformans' J1 and *Dehalobacterium formicoaceticum* J2.

**Table S5.** Genes involved in anaerobic *de novo* cobamide biosynthesis identified in the genomes of 'Ca. Dehalobacter formatiformans' J1 and *Dehalobacterium formicoaceticum* J2.
